# Overexpression of diglucosyldiacylglycerol synthase leads to daptomycin resistance in *Bacillus subtilis*

**DOI:** 10.1101/2024.05.10.593546

**Authors:** Ryogo Yamamoto, Kazuya Ishikawa, Yusuke Miyoshi, Kazuyuki Furuta, Shin-Ichi Miyoshi, Chikara Kaito

## Abstract

The lipopeptide antibiotic daptomycin exhibits bactericidal activity against gram-positive bacteria by forming a complex with phosphatidylglycerol (PG) and lipid II in the cell membrane, causing membrane perforation. With the emergence of daptomycin-resistant bacteria, understanding the mechanism of bacterial resistance to daptomycin has gained great importance. In the present study, we found that overexpression of *ugtP*, which encodes a diglucosyldiacylglycerol synthase, resulted in daptomycin resistance in *Bacillus subtilis*. The amount of diglucosyldiacylglycerol (Glc_2_DAG) increased in the *ugtP*-overexpressed strain, whereas the amounts of the acidic phospholipids cardiolipin (CL) and PG and the basic phospholipid lysylphosphatidylglycerol (Lys-PG) decreased. Moreover, *ugtP* overexpression did not alter the cell surface charge or the susceptibility to the cationic antimicrobial peptide nisin or the cationic surfactant hexadecyltrimethylammonium bromide (CTAB). Furthermore, under daptomycin exposure, we obtained daptomycin-resistant mutants that carried *ugtP* mutations and showed increased amounts of Glc_2_DAG and more than 4-fold increase in the MIC of daptomycin. These results suggest that an increase in the amount of Glc_2_DAG changes the phospholipid composition and leads to daptomycin resistance without altering the bacterial cell surface charge in *B. subtilis*.

**Importance:** Daptomycin is one of the last-resort drugs for the treatment of methicillin-resistant *Staphylococcus aureus* (MRSA) infections, and the emergence of daptomycin-resistant bacteria has become a problem. Understanding the mechanism of daptomycin resistance is important for establishing clinical countermeasures against daptomycin-resistant bacteria. In the present study, we found that overexpression of *ugtP*, which encodes diglucosyldiacylglycerol synthase, causes daptomycin resistance in *B. subtilis*, a model organism for gram-positive bacteria. Overexpression of *ugtP* increased diglucosyldiacylglycerol levels, resulting in altered phospholipid composition and daptomycin resistance. These findings are important for establishing clinical strategies against daptomycin-resistant bacteria, including their detection.

## Introduction

The lipopeptide antibiotic daptomycin is used to treat infective endocarditis and deep-skin infections caused by methicillin-resistant *Staphylococcus aureus* (MRSA). In the bacterial cell membrane, daptomycin forms a complex with phosphatidylglycerol (PG) and lipid II in a Ca^2+^-dependent manner, causing membrane perforation in gram-positive bacteria (1). With the growing emergence of daptomycin-resistant gram-positive bacteria, a better understanding of daptomycin-resistance mechanisms has become essential. In *S. aureus*, mutations in the *mprF* gene, which are involved in the lysylation of PG, cause daptomycin resistance with or without effects on the bacterial cell surface charge (2–6). In *Enterococcus faecalis*, daptomycin resistance is caused by mutations in *liaF*, which is involved in the stress-sensing response of the cell membrane, or in *gdpD*, which is involved in phospholipid metabolism (7). In *B. subtilis*, a mutation in the *pgsA* gene, which synthesizes PG, results in daptomycin resistance by reducing the amount of PG produced (8).

UgtP is an enzyme synthesizing diglucosyldiacylglycerol (Glc_2_DAG), a glyceroglycolipid, from UDP-glucose and diacylglycerol (DAG) (9). Glc_2_DAG is transformed into GP-Glc_2_DAG by reaction with PG (10) and functions as an anchor for lipoteichoic acid (LTA) in gram-positive bacteria (11,12). The *ugtP* gene is involved in various bacterial phenotypes; *the B. subtilis ugtP-*deletion mutant does not produce Glc_2_DAG, has an elongated LTA, and shows reduced swarming activity and increased expression of sigma factors that respond to the external environment (13–15). An *S. aureus* deletion mutant of *ypfP*, a homolog of *ugtP*, showed reduced amounts of LTA (16), higher levels of autolysis (16,17), decreased colony spreading (18), and decreased secretion of leukocidin (LukAB) (19). In invasive Group A *Streptococcus*, Glc_2_DAG plays a role in evading attacks from immune cells (20).

The *ugtP* gene is also involved in the susceptibility to several antimicrobial substances. *Bacillus subtilis ugtP*-deletion mutants are susceptible to sublancin, a cationic antimicrobial peptide (14). The *ugtP*-deletion mutants of MRSA with altered cell size and cell wall integrity are susceptible to β-lactam antibiotics and cell wall hydratases (21). However, the relationship between *ugtP* and daptomycin resistance has not been investigated. In the present study, we aimed to determine the genetic factors involved in daptomycin resistance in *B. subtilis*, a model gram-positive bacterium, and demonstrated that the overexpression of *ugtP* in *B. subtilis* leads to daptomycin resistance.

## Results

### Overexpression of Glc_2_DAG synthase in *B. subtilis* leads to daptomycin resistance

Using the *B. subtilis* gene knockout library, we searched for a gene knockout strain that grew on Luria-Bertani (LB) agar plates containing 4 µg/mL of daptomycin (22). The *metA*-deletion mutant showed resistance to daptomycin (**Fig. 1A**). In the absence of daptomycin, the *metA*-deletion mutant showed colony formation comparable to that of the wild-type strain on LB agar plates, but showed slightly delayed growth in liquid LB (**Fig. 1A, Fig. S1**). To confirm that the lack of *metA* leads to daptomycin resistance, we performed a complementation experiment. The *metA* gene was introduced into the *amyE* locus of *B. subtilis* to construct a *metA*-complemented strain. Introducing the *metA* gene into the *metA*-deletion mutant did not alter daptomycin sensitivity (**Fig. S2**). Because *ugtP* is located downstream of the *metA* locus (**Fig. 1B**), we hypothesized that the daptomycin resistance of the *metA*-deletion mutant was due to the polar effect of the erythromycin resistance cassette (*ermC*), an antibiotic resistance marker, in the *metA* locus. Therefore, we constructed a *metA* markerless deletion mutant by removing *ermC* from the *metA*-deletion mutant (**Fig. 1B**). As expected, the *metA* markerless deletion mutant did not show daptomycin resistance (**Fig. 1A**). Furthermore, overexpression of *ugtP* in the wild-type strain using an isopropyl-beta-D-thiogalactopyranosid (IPTG)-inducible promoter resulted in daptomycin resistance (**Fig. 1A**). In the absence of daptomycin, the *ugtP*-overexpressed strain showed colony formation and growth comparable to that of the vector-transformed strain (**Fig. 1A, Fig. S1**). These results suggest that increased expression of *ugtP* leads to daptomycin resistance.

**Figure 1.**
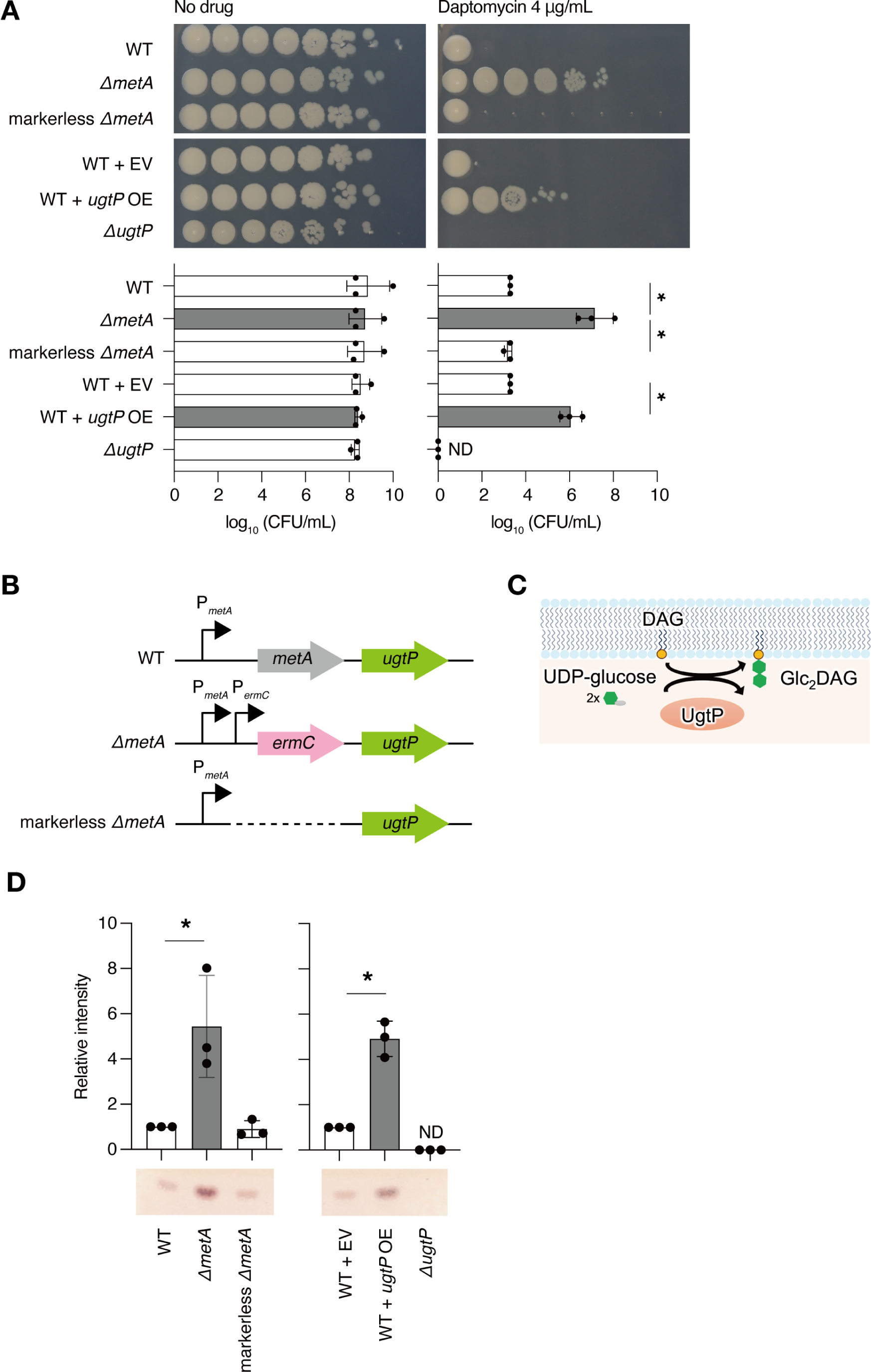
Increased amounts of Glc_2_DAG in *Bacillus subtilis* led to daptomycin resistance. A: Overnight cultures of the wild-type strain (WT), the *metA*-deletion mutant (Δ*metA*), the markerless *metA*-deletion mutant (markerless Δ*metA*), the wild-type strain transformed with empty vector (WT + EV), the wild-type strain transformed with *ugtP*-overexpressing plasmid (WT + *ugtP* OE), and the *ugtP*-deletion mutant strain (Δ*ugtP*) were serially diluted 10-fold, spotted onto Luria-Bertani (LB) agar plates containing 1 mM isopropyl-beta-D-thiogalactopyranosid supplemented with or without daptomycin 4 µg/mL, and incubated at 37°C overnight. The log_10_ CFU/mL value was calculated based on the number of colonies formed (graph). Data are presented as mean ± SD from three independent experiments. Statistical differences between the groups were analyzed using Tukey’s multiple-comparison test. *, P < 0.05. B: Schematic representation of the genome locus encompassing the *metA* and the *ugtP* genes in the WT, Δ*metA*, and markerless Δ*metA* strains. P*_metA_* and P*_ermC_* represent the promoter regions of the *metA* and erythromycin resistance genes (*ermC*), respectively. C: Schematic representation of diglucosyldiacylglycerol (Glc_2_DAG) synthesis in *B. subtilis*. In *B. subtilis*, UgtP synthesizes Glc_2_DAG from UDP-glucose and diacylglycerol (DAG). D: Total lipids of bacterial strains in A were extracted and analyzed using thin-layer chromatography (TLC). The signal intensity of Glc_2_DAG was measured. Data are expressed as mean ± SD from three independent experiments. Amounts relative to the WT (left graph) and WT + EV (right graph) are shown. Statistical differences between groups were analyzed using Student’s t-test. *, P < 0.05.

To investigate whether overexpression of *ugtP*, which encodes Glc_2_DAG synthase (**Fig. 1C**), increased the amount of Glc_2_DAG, total lipids were extracted from the bacteria and examined using thin-layer chromatography (TLC). The amounts of Glc_2_DAG in the *ugtP*-overexpressed strain and the *metA*-deletion mutant were greater than those in the vector-transformed and wild-type strains (**Fig. 1D**). The amount of Glc_2_DAG was not altered in the *metA* markerless deletion mutant in comparison with that in the wild-type strain (**Fig. 1D**). The *ugtP*-deletion mutant did not produce Glc_2_DAG (**Fig. 1D**) and was more susceptible to daptomycin than the wild-type strain (**Fig. 1A**). Since daptomycin resistance correlates with Glc_2_DAG accumulation, these results suggest that Glc_2_DAG synthesized by *ugtP* is involved in daptomycin resistance.

### Overexpression of Glc_2_DAG synthase results in decreased bactericidal activity of daptomycin

To analyze the effect of Glc_2_DAG on daptomycin activity in *B. subtilis* in more detail, we measured the number of dead bacteria stained with propidium iodide (PI) using flow cytometry. Bacterial cells with holes in their membranes due to daptomycin activity were expected to be stained with PI. After daptomycin exposure, the percentage of PI-positive cells in the *ugtP*-overexpressed strain was half that in the vector-transformed strain (**Fig. 2A, 2B**). These results suggest that the bactericidal activity of daptomycin was reduced in the *ugtP*-overexpressed strain.

**Figure 2.**
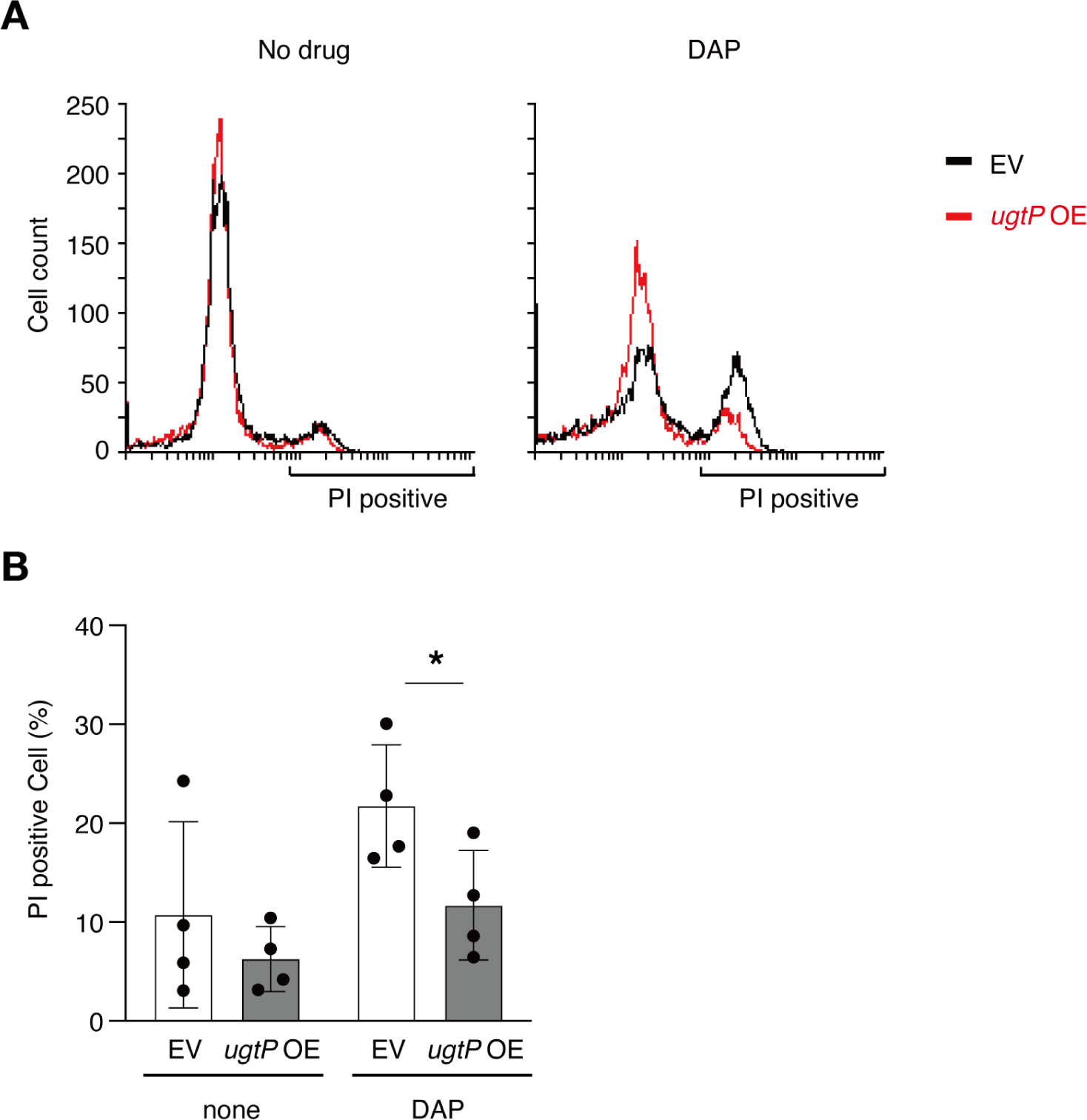
UgtP overexpression leads to reduced bactericidal ability of daptomycin. A: Logarithmically growing cells of the wild-type strain transformed with empty vector (EV) and the wild-type strain transformed with *ugtP*-overexpressing plasmid (*ugtP* OE) were stained by propidium iodide (PI) after treatment with or without daptomycin 10 μg/mL for 5 min at 37°C. The vertical axis represents the number of cells and the horizontal axis represents the fluorescence intensity of PI. B: The percentage of PI-positive cells was measured in A. Data are expressed as mean ± SD from three independent experiments. The statistical significance of the differences was analyzed using Student’s t-test. *, P < 0.05.

### Overexpression of Glc_2_DAG synthase did not change the bacterial cell surface charge

Based on a study reporting that the *mprF-*overexpressed *B. subtilis* strain shows daptomycin resistance along with a decrease in the negative charge of the cell membrane (23), we examined whether the *ugtP*-overexpressed strain also showed an altered cell surface charge by using the binding assay of cytochrome c, which binds to the negatively charged cell surface. No significant difference in cytochrome c binding was observed between the vector-transfected and *ugtP-*overexpressed strains (**Fig. 3A**). Furthermore, the *ugtP-*overexpressed strain and the vector-transformed strain showed similar sensitivity to the cationic antimicrobial agents nisin and hexadecyltrimethylammonium bromide (CTAB) (**Fig. 3B**). These results suggest that the daptomycin resistance caused by *ugtP* overexpression is not attributable to alterations in the cell surface charge.

**Figure 3.**
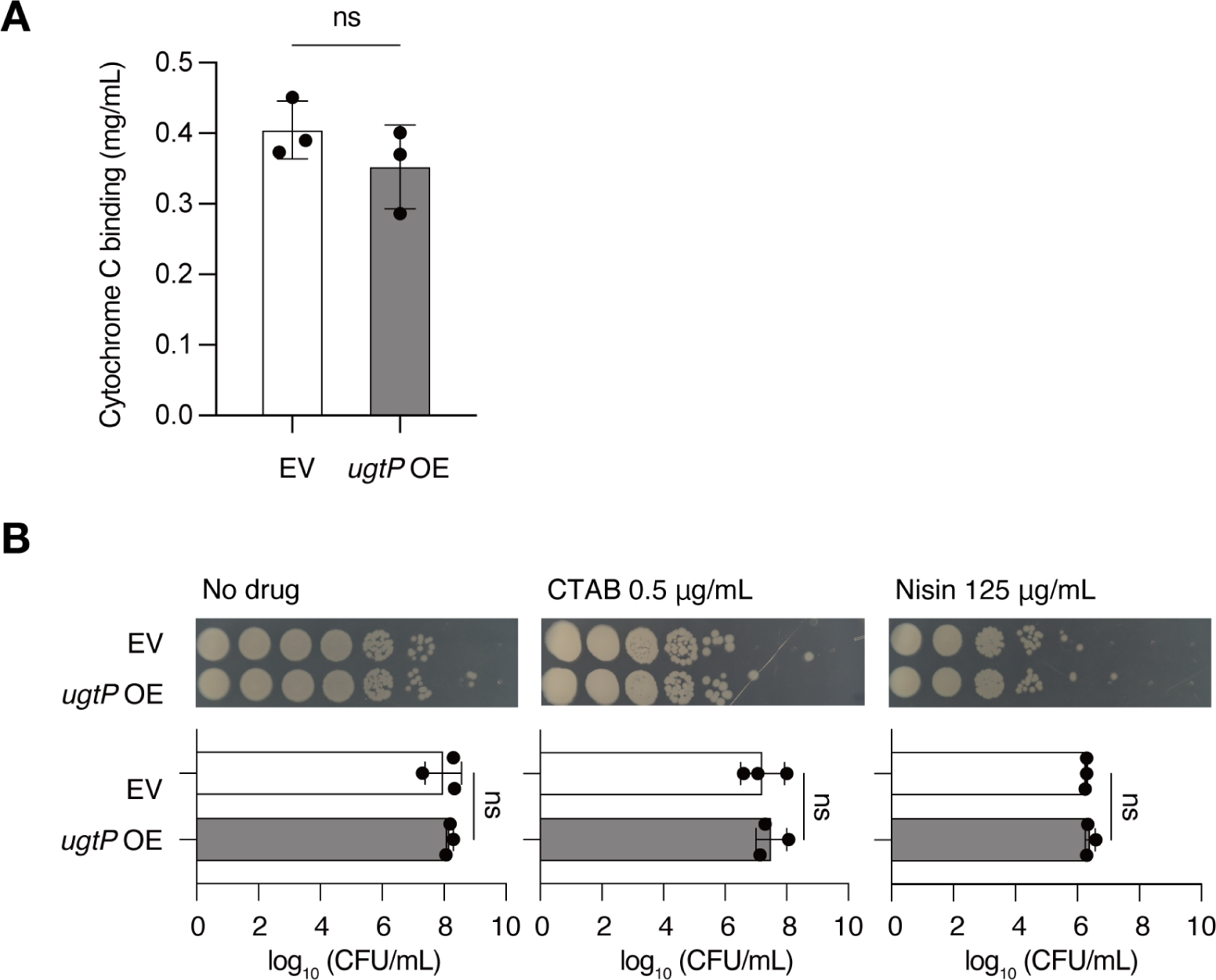
The *ugtP*-overexpressed strain does not show surface charge alterations. A: Surface charge of *B. subtilis* was evaluated by cytochrome c binding. Overnight cultures of the wild-type strain transformed with an empty vector (EV) and the wild-type strain transformed with *ugtP*-overexpressing plasmid (*ugtP* OE) were subjected to a cytochrome c binding assay. Data are presented as mean ± SD from three independent experiments. Statistical differences between groups were analyzed using Student’s t-test. ns, not significant. B: Overnight cultures of the wild-type strain transformed with EV and the wild-type strain transformed with *ugtP* OE were serially diluted 10-fold, spotted on Luria-Bertani (LB) agar plates supplemented with or without 0.5 µg/mL of hexadecyltrimethylammonium bromide or 125 µg/mL of nisin, and incubated at 37°C overnight. Log_10_ CFU/mL was calculated from the number of colonies formed (graph below). Data are presented as mean ± SD from three independent experiments. The statistical significance of differences between groups was analyzed using Student’s t-test. ns, not significant.

### Overexpression of Glc_2_DAG synthase alters the phospholipid composition of the cell membrane

On the basis of previous studies showing that decreased amounts of PG lead to daptomycin resistance in *B. subtilis* and *S. aureus* (8,24), we compared the amounts of phospholipids in the vector-transformed and *ugtP*-overexpressed strains by TLC. The amounts of the major phospholipids of *B. subtilis*, namely, cardiolipin (CL), PG, and lysylphosphatidylglycerol (Lys-PG), were determined by comparing the Rf values with those reported previously (25). Compared to the vector-transformed strain, the *ugtP*-overexpressed strain showed lower amounts of CL, PG, and Lys-PG (**Fig. 4A, 4B**). The amount of phosphatidylethanolamine (PE) was not significantly different between the *ugtP*-overexpressed and vector-transformed strains (**Fig. 4A, 4B**). These results suggest that *ugtP* overexpression decreases the levels of CL, PG, and Lys-PG.

**Figure 4.**
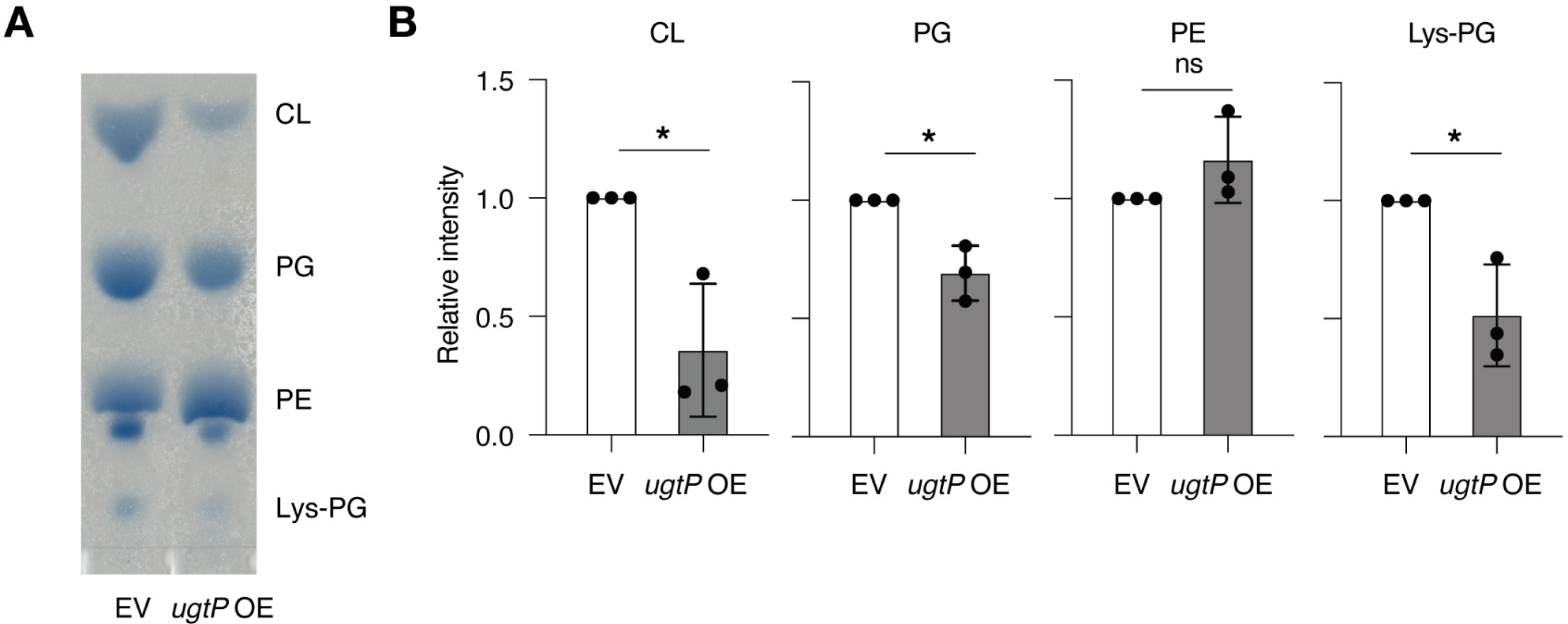
UgtP overexpression alters phospholipid composition. A: Total lipids were extracted from overnight cultures of the wild-type strain transformed with an empty vector (EV) and the wild-type strain transformed with *ugtP*-overexpressing plasmid (*ugtP* OE) and analyzed by thin-layer chromatography. CL, cardiolipin; PG, phosphatidylglycerol; PE, phosphatidylethanolamine; Lys-PG, lysylphosphatidylglycerol. B: Signal intensities of CL, PG, PE, and Lys-PG were measured in A. Data are shown as mean ± SD from three independent experiments. Relative amounts of phospholipids in the EV strains are shown. The statistical significance of differences between groups was analyzed using Student’s t-test. * P < 0.05, significant difference; ns, not significant.

### The emergence of daptomycin-resistant strains producing a large amount of Glc_2_DAG in an experimental evolution

To investigate whether mutants with increased Glc_2_DAG accumulation emerged after daptomycin exposure, we performed serial passaging in the presence of daptomycin. Six lineages were independently cultured for 15-26 passages in the presence of daptomycin, and four strains ( A, B, D, and E) with more than 4-fold higher MIC than that of the wild-type strain were obtained (**Fig. 5A**). TLC analysis revealed that the amount of Glc_2_DAG was increased in two of the four daptomycin-resistant strains (**Fig. 5B**). To determine whether these daptomycin-resistant mutants carried mutations in the *ugtP* gene, the *ugtP* locus was amplified by PCR and sequenced. In strain E, the PCR product was larger than that of the wild-type strain (**Fig. 5C**), and sequence analysis revealed that a partial region encompassing the *ugtP* promoter region was duplicated (**Fig. 5D**). In strain D, guanine at position 226 was replaced by cytosine, resulting in the exchange of valine-76 of UgtP for leucine (**Fig. 5D**). In the other two daptomycin-resistant strains, which did not show an increase in the amount of Glc_2_DAG, no mutations were observed in the *ugtP* gene. These results indicated that daptomycin exposure results in an increase in the amount of Glc_2_DAG in daptomycin-resistant strains, which is caused by mutations in the *ugtP* gene.

**Figure 5.**
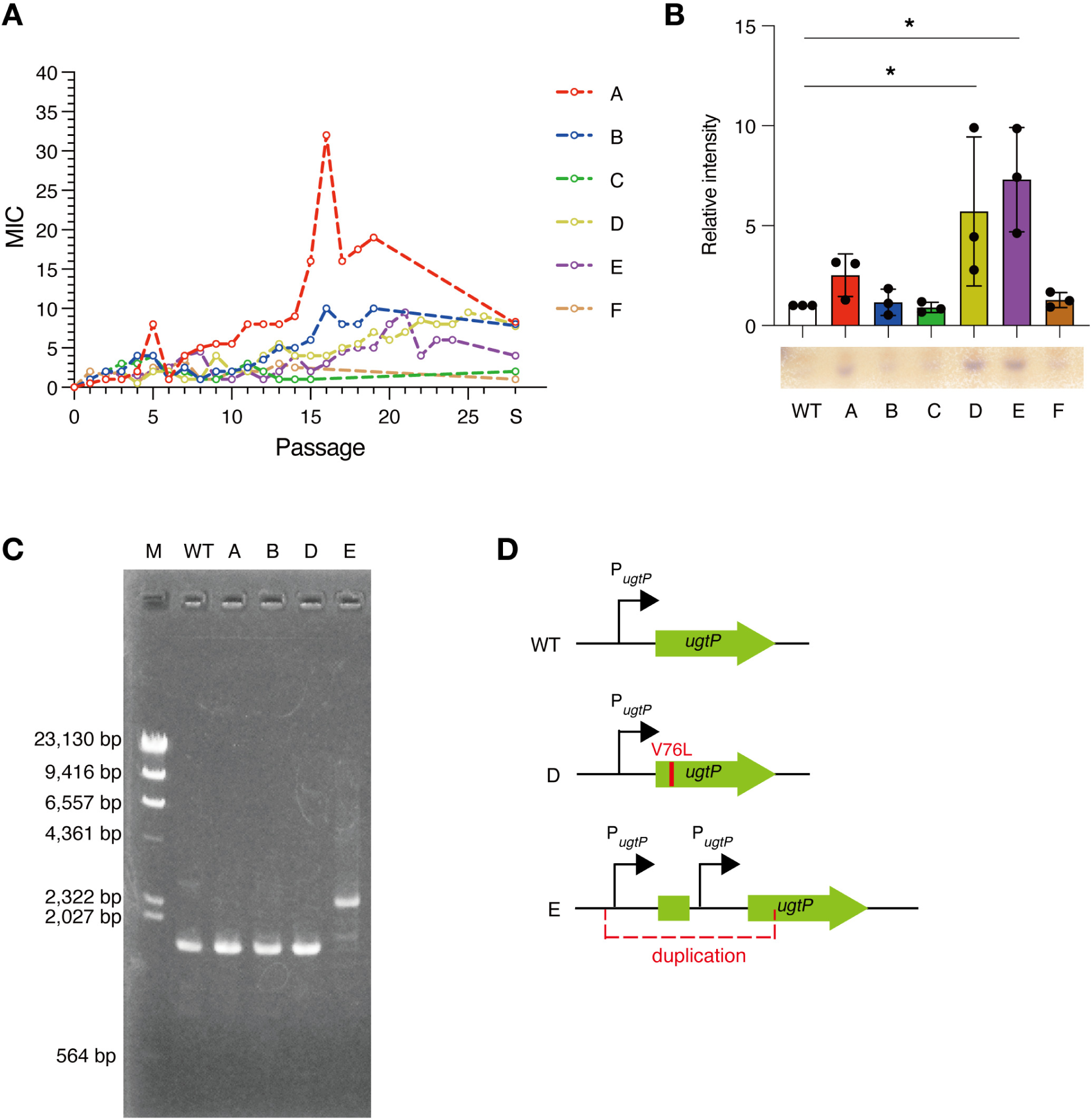
Daptomycin-resistant strains isolated in an experimental evolution have *ugtP* mutations and increased amounts of diglucosyldiacylglycerol. A: Six lineages of *B. subtilis* (A–F) were serially passaged in daptomycin-containing medium and their MICs were measured. The horizontal axis represents the number of passages, and the vertical axis represents the daptomycin MIC. Finally, a single colony was isolated from the passaged cultures, named strains A–F, for which the MICs are shown at S on the horizontal axis. B: Total lipid content was extracted from six strains (A–F), and the amount of diglucosyldiacylglycerol was measured by thin-layer chromatography. Data are shown as mean ± SD from three independent experiments. The relative amounts of diglucosyldiacylglycerol to those in the wild-type strain (WT) are shown. The significance of differences between groups was analyzed using Dunnett’s multiple-comparison test. *, P < 0.05. C: The *ugtP* region was amplified by PCR and subjected to DNA electrophoresis. D: Schematic representation of the *ugtP* locus in the wild-type, D, and E strains.

## Discussion

In this study, *ugtP* overexpression led to daptomycin resistance in *B. subtilis*. To our knowledge, this is the first report describing the relationship between increased amounts of Glc*_2_*DAG and daptomycin resistance in *B. subtilis*.

Overexpression of *ugtP* in *B. subtilis* increased the amount of Glc_2_DAG and altered the amount of phospholipids (**Fig. 1, 4)**. Because Glc_2_DAG reacts with the phosphoglycerol group of PG to form GP-Glc_2_DAG (26), the increased Glc_2_DAG levels in the *ugtP*-overexpressed strain would lead to a decrease in the amount of PG (**Fig. 6**). This may have decreased the efficiency of complex formation between daptomycin and PG and resulted in daptomycin resistance (**Fig. 6**), because daptomycin forms a complex with PG and causes membrane perforation (1). In contrast, CL and Lys-PG levels decreased in the *ugtP*-overexpressed strain. The decrease in these phospholipids may also be involved in daptomycin resistance, because CL is a dimer of PG (27) and Lys-PG is a modified PG (**Fig. 6**). While the relative amount of Glc_2_DAG to total lipids has been reported to be 10% in exponentially growing *B. subtilis* cells (28), this study revealed that the *ugtP*-overexpressed strain produced 5-fold more Glc_2_DAG than the vector-transformed strain (**Fig. 5**). Thus, the *ugtP*-overexpressed strain presumably produces a large amount of Glc_2_DAG in the total lipid content, which may directly affect the action of daptomycin. Further investigation is needed to examine the effects of changes in lipid composition on daptomycin activity.

**Figure 6.**
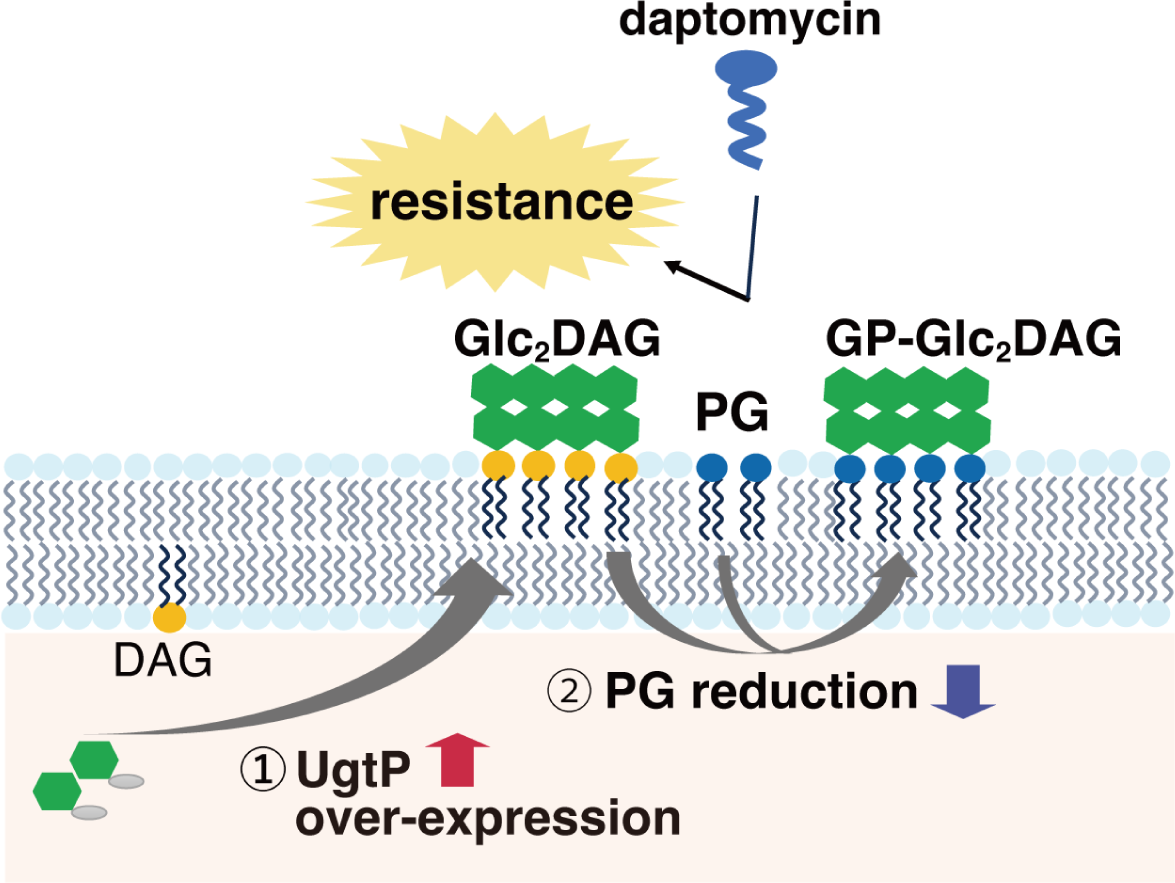
Model in which *ugtP* overexpression leads to daptomycin resistance. UgtP overexpression increases diglucosyldiacylglycerol levels (1) and decreases phosphatidylglycerol (PG) levels (2). A decrease in PG levels may decrease the efficiency of the daptomycin-PG complex formation.

Daptomycin resistance is not explained as electric repulsion between the bacterial cell surface and daptomycin in *S. aureus* (24). In the present study, the cell surface charge of the *ugtP*-overexpressed *B. subtilis* strain was not altered, although the amounts of the acidic phospholipids CL and PG decreased (**Fig. 3, 4)**. The decreased amount of the basic phospholipid Lys-PG may have counteracted the effect of the decreased amounts of acidic phospholipids in the *ugtP*-overexpressed strain. In the present study, the *ugtP*-overexpressed strain showed similar sensitivity to cationic antimicrobial substances such as nisin and CTAB as the vector-transformed strain. Therefore, the daptomycin resistance of the *ugtP-*overexpressed *B. subtilis* strain was not due to alterations in the cell surface charge.

This study also revealed elevated Glc_2_DAG levels and daptomycin resistance in *B. subtilis* strains serially passaged in a daptomycin-containing medium. Sequencing analysis identified two independent spontaneous mutations in *ugtP*; duplication of the *ugtP* promoter and an amino acid substitution of UgtP, which could increase *ugtP* transcription and UgtP activity/stability, respectively, resulting in an increase in the amount of Glc_2_DAG. Therefore, the spontaneous mutations in the *ugtP* gene cause resistance to daptomycin. In *E. faecalis* and *S. aureus*, the amount of Glc_2_DAG was shown to be increased in daptomycin-resistant strains (29). Thus, based on the findings of this study, the increased amount of Glc_2_DAG may be a major factor contributing to daptomycin resistance in gram-positive bacteria. Future studies should aim to investigate the involvement of *ugtP* homologues in bacteria other than *B. subtilis* and the molecular mechanisms by which increased amounts of Glc_2_DAG cause daptomycin resistance.

## Materials and Methods

### Strains and culture conditions

*B. subtilis* 168 *trpC2* and the mutant strains were cultured on LB agar plates or in LB liquid medium under aerobic conditions at 37°C. The *B. subtilis* gene deletion mutants carrying *ermC* were similarly cultured on LB agar plates or in LB liquid medium containing 1 µg/mL erythromycin. To overexpress UgtP, bacterial strains were cultured on LB agar plates or in LB liquid medium containing 1 mM IPTG. Details of the bacterial strains and plasmids used are listed in Table 1.

**Table 1.** List of bacterial strains and plasmids used.

| Strain or plasmid | Genotype or characteristics | Source or reference |
| --- | --- | --- |
| Strains |  |  |
| *Bacillus subtilis* |  |  |
| 168 | *trpC2* | BGSC |
| BKE13680 | *trpC2*, Δ*motB::erm*, *Erm<sup>r</sup>* | [22] |
| 168 (BKE) | *trpC2*, originated from BKE13680 | This study |
| BKE21910 | *trpC2*, Δ*metA::erm*, *Erm<sup>r</sup>* | [22] |
| BKE21920 | *trpC2*, Δ*ugtP::erm*, *Erm<sup>r</sup>* | [22] |
| BKE21910ML | *trpC2*, Δ*metA::markerless* | This study |
| *Escherichia coli* |  |  |
| JM109 | Host strain for cloning | Takara Bio |
| Plasmids |  |  |
| pDR110 | An integration vector, *Amp<sup>r</sup>*, *Spc<sup>r</sup>* | BGSC [22] |
| pDR110-*metA* | pDR110 with *metA*, *Amp<sup>r</sup>*, *Spc<sup>r</sup>* | This study |
| pDR110-*ugtP* | pDR110 with *ugtP*, *Amp<sup>r</sup>*, *Spc<sup>r</sup>* | This study |
| pDR244 | Cre recombinase-expressing plasmid, *Amp<sup>r</sup>*, *Spc<sup>r</sup>* | BGSC [22] |

### Screening of daptomycin-resistant mutants

The BKE library (22) was cultured in 96-well microplates at 37°C. The cultures were spotted onto agar plates containing 4 µg/mL of daptomycin using a replicator and incubated at 37°C overnight. Mutant strains that formed colonies were identified. The identified strains were re-streaked onto daptomycin-containing plates and their resistance was confirmed.

### Genetic manipulations

#### (1) Construction of the wild-type strain

Since the colony shapes of the gene deletion mutants in the BKE library and the 168 *trpC2* strain from BGSC were different, we assumed that the genetic background differed between the parent strain of the BKE library and the 168 *trpC2* strain from BGSC. To compare the phenotypes of the BKE library mutant strains with those of the wild-type strain, we constructed a wild-type strain from the *motB*-deletion mutant (BKE13680) in the BKE library by repairing the *motB* gene. The *motB*-deletion mutant was transformed with DNA fragments containing the *motB* gene, cultured, and 2 μL was inoculated into the center of 0.3% LB agar plates. After overnight incubation at 37℃, the peripheral region of the colony was picked with a toothpick, and a single colony was isolated. The isolated strain was again inoculated into the center of 0.3% LB agar plates, and recovery of the swimming activity was confirmed. Correct repair of the *motB* locus was confirmed by PCR using specific oligonucleotide primers (**Table 2**). The colony shape was similar to that of the BKE library mutants. This strain was used as the wild-type strain in the present study.

**Table 2.**
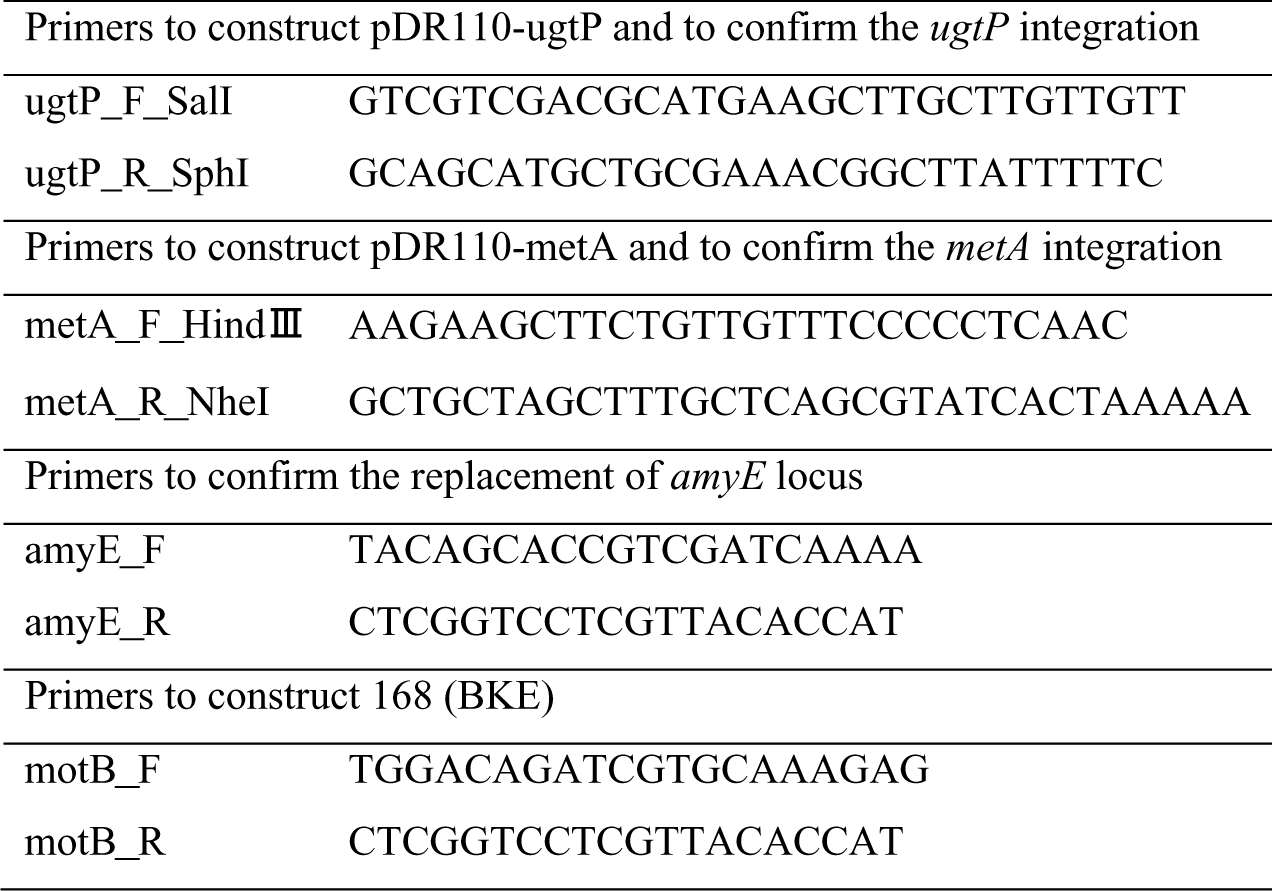
Primers used in this study.

#### (2) Construction of gene deletion mutants

Genomic DNA from the *metA* or *ugtP*-deletion mutants in the BKE library was extracted using the QIAamp DNA Blood Mini Kit (Qiagen). The extracted genomic DNA was used to transform the *B. subtilis* wild-type strain, which was spread on agar plates containing erythromycin (1 µg/mL) and incubated overnight at 37°C. Erythromycin-resistant colonies were subjected to colony PCR to confirm the presence of the *metA* or *ugtP* deletions.

#### (3) Construction of markerless *metA*-deletion mutant

The *metA*-deletion mutant was transformed with pDR244 (22) and passaged twice on agar plates containing spectinomycin (100 µg/mL) at 30°C. The colonies were then streaked onto LB agar medium without drugs and incubated overnight at 43°C. The growing colonies were then streaked onto drug-free LB agar plates and incubated overnight at 43°C. The resulting colonies were streaked onto agar plates containing erythromycin (100 µg/mL) or spectinomycin (100 µg/mL), and confirmed to be sensitive to these antibiotics. PCR was performed to confirm the loss of erythromycin resistance markers from the *metA* gene region.

#### (4) Construction of *ugtP*-overexpressed strain

DNA fragments containing the *ugtP* gene were amplified by PCR using oligonucleotide primers (**Table 2**) and genomic DNA of the wild-type strain as a template. The amplified DNA fragment was inserted into the Sal1 and Sph1 regions of pDR110 to obtain pDR110-*ugtP*. The plasmid was used to transform the *B. subtilis* wild-type strain and cultured on LB agar plates containing spectinomycin (100 µg/mL). Correct insertion of *ugtP* into the *amyE* locus was confirmed by PCR.

### Evaluation of bacterial resistance to antibiotics

To evaluate antibiotic resistance, the autoclaved LB agar medium and antibiotic solution were mixed and poured into square petri dishes (Eiken Chemical, Tokyo, Japan). Overnight cultures of the bacterial strains were serially diluted 10-fold in 96-well microplates, and 5 µL of the diluted bacterial solution was spotted on the LB agar plates containing antibiotics and incubated at 37°C. The plates were photographed using a digital camera, and the number of colonies was counted according to a previously described method (30).

### Measurement of bacterial growth curve

Overnight cultures (2 µL) of *B. subtilis* strains were inoculated into 200 µL of LB medium containing 1 mM IPTG in a 96-well microplate and incubated at 37°C for 7 h with shaking. The OD_595_ was measured using a microplate reader (Muktiskan FC, Thermo Fisher Scientific).

### Experimental evolution of daptomycin-resistant *B. subtilis* strains

*B. subtilis* colonies were inoculated into 5 mL of LB medium supplemented with 5 µL of ethyl methanesulfonate (EMS), and cultured at 37°C overnight. The EMS-treated culture (2 µL) was inoculated into LB medium (200 µL) containing 0.125 mM calcium chloride and various concentrations of daptomycin (0.5-µg/mL increments), and incubated at 37°C without shaking overnight. Bacterial growth in the well containing the highest concentration of daptomycin was inoculated again into LB medium (200 µL) containing 0.125 mM calcium chloride and various concentrations of daptomycin. The culture process in the presence of daptomycin was repeated 15–26 times, and four resistant strains with a higher MIC than that of the wild-type strain were isolated. The *ugtP* region of the resistant strains was amplified using PCR and subjected to Sanger sequencing.

### Lipid extraction and TLC

Total lipid extraction and TLC analysis of Glc_2_DAG were performed as described previously with minor modifications (31). Overnight cultures (1 mL) of *B. subtilis* strains were inoculated in 100 mL of LB medium in a 0.5-L flask, cultured aerobically at 37°C for 4 h 30 min, after which 40 mL of the bacterial culture was centrifuged at 10,400 × *g* for 10 min at 4°C. The bacterial pellet was suspended in 1 mL of MilliQ water, and lipids were extracted using the Bligh and Dyer method (32). The lipid fraction was evaporated in a centrifugal evaporator, and the lipids were dissolved in 200 µL of chloroform:methanol (1:1 v/v). The samples were spotted onto a TLC silica gel 60 plate (Merck). The plate was developed in chloroform:methanol:water (65:25:4 v/v), sprayed with a coloring agent (10.5 mL of 15% 1-naphthol in ethanol, 6.5 mL of sulfuric acid, and 4 mL of water), and then heated at 110°C. For phospholipid analysis, a previously described method was used with minor modifications (25,33). Total lipids were extracted from 500 mL of the bacterial culture using the method described above. Before spotting the samples, TLC plates were immersed in 1.8% boric acid in ethanol for 2 min, dried for 15 min, and heated at 100°C for 15 min. After spotting the sample, the TLC plates were developed in chloroform:methanol:water:ammonia (56:35:2.8:1, v/v/v/v). Molybdenum blue spray reagent (MERCK) was used to visualize the phospholipids.

### Evaluation of the bactericidal activity of daptomycin

The bactericidal activity of daptomycin was evaluated as described previously (34). Logarithmically growing *B. subtills* cells were centrifuged at 4000 × *g* for 5 min, and the bacterial pellet was suspended in 1 mL of phosphate-buffered saline (PBS), diluted 10-fold, treated with 10 µg/mL of daptomycin at 37°C for 5 min, and then centrifuged at 4000 × *g* for 5 min. The bacterial pellet was suspended in 1 mL of PBS, mixed with 1.33 µg/mL PI, and incubated at 37°C for 15 min in the dark. The percentage of bacteria that took up PI was measured by flow cytometry.

### Cytochrome c binding assay

The cytochrome c binding assay was performed as described previously (35). Overnight cultures of *B. subtills* strains were centrifuged at 2500 × *g* for 10 min. The bacterial pellet was washed twice with buffer (20 mM MOPS, 5 mM sodium citrate, and 1 mM EDTA, pH 7.0). The cells were suspended in the same buffer and adjusted to an OD_600_ of 7. Next, 0.5 mg/mL cytochrome c was added to the cells and incubated at 37°C for 15 min. The cells were centrifuged, and the OD_530_ of the supernatant was measured. The cytochrome c concentration in the supernatant was calculated using a standard curve.

### Statistical analysis

All statistical analyses were performed using Prism software (version 9.4.1; GraphPad Software).

## Acknowledgement

This study was supported by Japan Society for the Promotion of Science (JSPS) Grants-in-Aid for Scientific Research (Grants 22K14892, 23K24131, 23K06130, 24K01760) and the Program of the Japan Initiative for Global Research Network on Infectious Diseases (J-GRID), JP23wm0125004, from Ministry of Education, Culture, Sports, Science and Technology in Japan (MEXT), and Japan Agency for Medical Research and Development (AMED). This study was also supported by the Takeda Science Foundation (to C.K.), the Ichiro Kanehara Foundation (to C.K.), the Ryobi Teien Memory Foundation (to K.I. and C.K.). We thank the National BioResource Project-B. subtilis (National Institute of Genetics, Japan) for providing the B. subtilis BKE library and Bacillus Genetic Stock Center (BGSC) for providing the B. subtilis 168 trpC2 and B. subtilis plasmids.

---

**Figure S1.** Growth curves of the wild-type, Δ*metA*, and *ugtP-*overexpressing strains Growth curves of the wild-type, Δ*metA*, vector-transformed, and *ugtP-*overexpressing strains were examined. Data are presented as means ± standard error (n = 3).

**Figure S2.** Daptomycin resistance is not canceled by *metA* complementation of Δ*metA* Overnight cultures of the wild-type strain (WT), the *metA*-deletion mutant (Δ*metA*), and the Δ*metA* complementation strain with *metA* at the *amyE* locus of Δ*metA* (Δ*metA*/*spec*-P_spank_-*metA*) were serially diluted 10-fold, spotted onto Luria-Bertani agar plates containing 1 mM isopropyl-beta-D-thiogalactopyranosid supplemented with or without daptomycin 4 µg/mL, and incubated at 37°C overnight.

